# Ethanol Experience Enhances Glutamatergic Ventral Hippocampal Inputs to D1 Receptor-Expressing Medium Spiny Neurons in the Nucleus Accumbens Shell

**DOI:** 10.1101/471011

**Authors:** Daniel M. Kircher, Heather Aziz, Regina A. Mangieri, Richard A. Morrisett

## Abstract

Nucleus accumbens dopamine D1 receptor-expressing medium spiny neurons (D1-MSNs) have been implicated in the formation of dependence to many drugs of abuse including alcohol. Previous studies have revealed that acute alcohol exposure suppresses glutamatergic signaling within the accumbens and repeated alcohol exposure enhances glutamatergic signaling. D1-MSNs receive glutamatergic input from several brain regions and it is not currently known how individual inputs onto D1-MSNs are altered by alcohol experience. To Address this, we used virally mediated expression of Channelrhodopsin (ChR2) in ventral hippocampal (vHipp) glutamate neurons to selectively activate vHipp to D1-MSN synapses and compared synaptic adaptations in response to low and high alcohol experience *in vitro* and *in vivo*. Alcohol experience enhanced glutamatergic activity and abolished long-term depression (LTD) at ventral hippocampal (vHipp) to D1-MSN synapses. Following chronic alcohol experience GluA2-lacking AMPA receptors, which are Ca-permeable, were inserted into vHipp to D1-MSN synapses. These alcohol-induced adaptations of glutamatergic signaling occurred at lower levels of exposure than previously reported. The loss of LTD expression and enhancement in glutamatergic signaling from the vHipp to D1-MSNs in the nucleus accumbens may play a critical role in the formation of alcohol dependence and enhancements in ethanol consumption. Reversal of alcohol-induced insertion of Ca-permeable AMPA receptors and enhancement of glutamatergic activity at vHipp to D1-MSNs presents potential targets for intervention during early exposure to alcohol.

**SIGNIFICANCE STATEMENT:** The work presented here is the first to elucidate how an individual glutamatergic input onto D1-MSNs of the accumbens shell (shNAc) are altered by repeated ethanol exposure. Our findings suggest that glutamatergic input from the ventral hippocampus (vHipp) onto D1-MSNs is enhanced following drinking in a two-bottle choice (2BC) paradigm and is further enhanced by chronic intermittent ethanol (CIE) vapor exposure which escalated volitional ethanol intake. A critical finding was the insertion of Ca-permeable AMPA receptors into vHipp-shNAc D1-MSN synapses following CIE exposure, and more importantly following ethanol consumption in the absence of vapor exposure. These findings suggest that enhancements of glutamatergic input from the vHipp and insertion of Ca-permeable AMPARs play a role in the formation of ethanol dependence.

## INTRODUCTION

Chronic exposure to drugs of abuse and alcohol induces several forms of plasticity within the nucleus accumbens (NAc) (Lüscher and Malenka, 2011; McCool, 2011). For example, several groups have shown that chronic exposure to psychostimulants, heroin, or alcohol alters the expression of NMDA receptor (NMDAR)-dependent long-term depression (LTD) in the NAc (Thomas et al., 2001; Martin et al., 2006; Mao et al., 2009; Kasanetz et al., 2010; Shen and Kalivas, 2012; Abrahao et al., 2013; Jeanes et al., 2014; Spiga et al., 2014; Renteria et al., 2017, 2018). Alterations in accumbal synaptic plasticity likely play a critical role in the formation of dependence on various drugs of abuse including alcohol.

Previous work in our lab demonstrated that chronic intermittent ethanol (CIE) vapor exposure enhances the excitability of dopamine D1 receptor-expressing medium spiny neurons (D1-MSNs) and increases NMDAR activity in D1-MSNs (Renteria et al., 2017). CIE reliably induces enhancements in volitional ethanol intake (Becker and Lopez, 2004; Jeanes et al., 2011; Renteria et al., 2017). Interestingly, CIE vapor exposure also disrupts glutamatergic synaptic plasticity in the NAc and does so in a manner that is dependent on the subtype of MSN and strain of mouse. Twenty-four hours after CIE vapor exposure, a plasticity induction protocol that normally produces NMDAR-dependent LTD of AMPAR-mediated excitatory postsynaptic currents (EPSCs) in D1-MSNs had no effect on AMPAR-EPSCs, when tested in mice of a Swiss Webster background strain (Jeanes et al., 2014). By contrast, when tested at the same time point post-CIE vapor, the same induction protocol produced long-term potentiation (LTP) of AMPAR-EPSCs in D1-MSNs from mice of a C57BL/6J background strain (Renteria et al., 2018). Differences in the effect of ethanol exposure on plasticity was likely due to C57BL/6J mice, but not Swiss Webster, readily consuming ethanol (Belknap et al., 1993). In other words, ethanol-induced alterations in glutamatergic synaptic plasticity (i.e. metaplasticity) within the D1-MSNs of the accumbens may be critically-related to the expression of ethanol drinking behavior, with enhanced glutamatergic activity within the NAc following CIE exposure leading to a high drinking phenotype compared to animals that do not exhibit such alterations.

To investigate this, Renteria et al. (2018) measured LTD in the NAc of C57BL/6J mice that exhibited escalated drinking following CIE exposure. The disruption in the nucleus accumbens shell (shNAc) D1-MSN plasticity persisted for up to two weeks and returned to air control levels three weeks after the last ethanol drinking session, mirroring the time course of escalated drinking observed following CIE exposure (Becker and Lopez, 2004; Renteria et al., 2018). The loss of LTD was also accompanied by an increase in the rectification index of AMPAR-mediated currents in D1-MSNs of the shNAc, 24 h after the last drinking session. The presence of rectification suggested synaptic insertion of GluA2 subunit-lacking AMPA receptors (AMPARs), which are Ca-permeable. Changes in rectification as well as insertion of Ca-permeable AMPARs into the synapse has been observed following withdrawal from cocaine administration (Conrad et al., 2008; Wolf and Tseng, 2012; Loweth et al., 2014). Insertion of Ca-permeable AMPARs into shNAc D1-MSN synapses following repeated drug exposure likely plays a role in relapse following periods of withdrawal as well as in the escalation of drug intake observed following repeated use and may play a role in the loss of LTD expression.

Recent studies with psychostimulants indicate that drug exposure produces selective effects on different glutamatergic inputs to the NAc (Pascoli et al., 2012, 2014; Stuber et al., 2012; Joffe and Grueter, 2016). Here, we designed a study to characterize neuroadaptations in a specific glutamatergic input to the shNAc with excessive ethanol consumption. The NAc receives major inputs from the hippocampal ventral subiculum (Groenewegen et al., 1987) as well as the ventral CA1 (Britt et al., 2012). Furthermore, the NAc requires hippocampal stimulation for depolarization and transition into an activated state (O’Donnell and Grace, 1995). Information from the hippocampus is involved in mediating drug-seeking behaviors (Fuchs et al., 2005; Riaz et al., 2017), with lesions altering Pavlovian conditioned approach behaviors (Fitzpatrick et al., 2016), that may be due to disrupting gating and outflow of NAc signals (French and Totterdell, 2002). Inactivation of the ventral CA1 region selectively inhibits cocaine seeking without disrupting food seeking or locomotor behavior (Rogers and See, 2007; Lasseter et al., 2010), suggesting that the ventral CA1 is likely a site where the persisting ‘memories that sustain addiction’ may be encoded (Nestler, 2001). Lastly, neurotransmission in vHipp-NAc circuitry has been implicated in triggering relapse in addiction (Kalivas et al., 2005). To our understanding no prior research has been conducted to examine the impact of ethanol exposure on vHipp to NAc shell circuitry. Therefore, we used optogenetics to selectively stimulate vHipp inputs to genetically identified D1-MSNs in the shNAc to determine if excessive alcohol consumption alters AMPAR-mediated signaling and disrupts NMDAR-LTD in this important glutamatergic projection to the shNAc.

## METHODS

All procedures were approved by the Institutional Animal Care and Use Committee of The University of Texas at Austin.

Mice were hemizygous BAC transgenic mice in which tdTomato fluorophore expression was driven by dopamine D1 receptor gene regulatory elements; B6.Cg-Tg(Drd1a-tdTomato)6Calak/J (Ade et al., 2011). *Drd1a*-tdTomato breeding pairs were previously obtained from The Jackson Laboratory (Stock No. 016204) and this line was maintained at the University of Texas at Austin by mating hemizygous mice with non-carrier *Drd1a*-tdTomato mice (or period backcrossing with C57BL/6J mice). All mice were kept in a temperature and humidity-controlled environment with a 12 h light/ 12 h dark cycle (lights off at 0930 h). All procedures were approved by the Institutional Animal Care and Use Committee of the University of Texas at Austin.

### Stereotaxic Injections

AAV2-CaMKIIa-hChR2(H134R)-eYFP or AAV2-CaMKIIa-eYFP (for control mice) produced at the University of North Carolina (Vector Core Facility) was injected into the ventral hippocampus (vHipp) of 5- to 6-week-old male mice. Anesthesia was induced at 3% and maintained at a range of 1.5-2% isoflurane (w/v) (Animal Health International). Mice were placed in a stereotaxic frame (Kopf) and bilateral craniotomies were performed using stereotaxic coordinates adapted from a mouse brain atlas (Franklin and Paxinos, n.d.) (anterior-posterior = −2.8; medial-lateral = ± 2.8; dorsal-ventral (from the surface of the brain) = −4.2). Injections of virus (0.55 μl per injection site) were made using a Nanoject II with pulled glass pipettes (Drummond Scientific Company), broken back to a tip diameter of 10-15 μm, at an infusion rate of ≈0.05 μl per second.

### Two Bottle Choice and Air/Ethanol Vapor Exposure

Two weeks after virus injections mice began 21 consecutive days of two bottle choice (2BC) drinking, a limited access (2 hours) paradigm used to measure volitional ethanol consumption (Griffin, 2014). Thirty minutes prior to the beginning of the dark cycle (lights out at 0930 hours) mice were given access to two bottles, one containing a 15% ethanol and tap water solution and the other containing tap water alone. Animals and bottles were weighed prior to 2BC and bottles were weighed again after two-hour access. The difference in bottle weights was used to calculate the dose of ethanol consumed as grams of ethanol (difference in ethanol bottle weights * 0.12) per kilogram animal body weight per 2 hours. The last 5 days of this drinking period was measured as baseline ethanol consumption.

Following this 21 days of drinking animals underwent chronic intermittent ethanol (CIE) vapor (or air) exposure (Becker and Lopez, 2004). Mice were exposed to ethanol vapor following an injection (i.p.) with a loading dose of ethanol (20% v/v, 1.5 g/kg) and pyrazole (68.1 mg/kg) suspended in 0.1M PBS. Air exposed animals received an injection of pyrazole but not ethanol. Ethanol was volatilized by bubbling air through a flask containing 95% ethanol at a rate of 3.5 liter/min. Ethanol vapor was then combined with a separate air stream to give a total flow rate of approximately 4 liters/min. Ethanol enriched air (or air alone for the Air group) was pumped into special mouse chamber units with airtight tops (Allentown Inc., Allentown NJ). Each CIE bout consisted of 4 consecutive days of exposure for 16 hours followed by 8 hours of withdrawal. Following each 16-hour exposure session blood ethanol concentrations (BEC) were analyzed from tail blood by samples by gas chromatograph (Bruker 430-GC, Fremont CA). Target BECs following each session were between 150-200 mg/dl (37-47 mM). BECs were maintained by adjusting the flow rate and the loading dose of ethanol and pyrazole. Additional details regarding BEC analysis can be found in Renteria et al., 2018. After the conclusion of each four-day bout of CIE or air exposure, animals were returned to their home cages for 72 hours of withdrawal. After withdrawal, 2BC drinking was conducted for 5 days to measure CIE-induced changes in ethanol consumption.

### Brain Slice Preparation

Parasagittal slices (230-240 μm thick) containing the shNAc were prepared from mice (> 13 weeks due to 2BC and CIE exposure following recovery from virus injection) using a Leica vibrating microtome. Mice were anesthetized by inhalation of isoflurane and the brains were rapidly removed and placed in 4°C oxygenated artificial cerebrospinal fluid (ACSF) containing the following in (mM): 210 sucrose, 26.2 NaHCO_3_, 1 NaH_2_PO_4_, 2.5 KCl, 11 dextrose 6 MgSO_4_, 2.5 CaCl_2_, bubbled with 95% O_2_/ 5% CO_2_. Slices were transferred to a non-sucrose based ACSF solution for incubation containing the following (in mM): 120 NaCl, 25 NaHCO3, 1.23 NaH_2_PO_4_, 3.3 KCl, 2.4 MgSO_4_, 1.8 CaCl_2_, 10 dextrose, were continuously bubbled with 95% O_2_/ 5% CO_2_; pH 7.4, 32°C, and were maintained in this solution for at least 60 minutes prior to recording.

### Patch Clamp Electrophysiology

Whole cell voltage clamp recordings were conducted in the medial shNAc. Cells were identified using a BX50 microscope (Olympus) mounted on a vibration isolation table. Recordings were made in ACSF containing (in mM): 120 NaCl, 25 NaHCO_3_, 1.23 NaH_2_PO_4_, 3.3 KCl, 0.9 Mg SO_4_, 2.0 CaCl_2_, and 10 dextrose, bubbled with 95% O_2_/ 5% CO_2_. ACSF was continuously perfused at a rate of 2.0 mL/min and maintained at a temperature of ≈ 32°C. Picrotoxin (50 μM) was included in the recording ACSF to block GABA_A_ receptor-mediated synaptic currents. The NMDAR antagonist APV (50 μM; Tocris) was added to the recording solution for Rectification Index (RI) recordings to isolate AMPAR mediated currents. Strontium chloride (SrCl_2_, 6 mM; Tocris) was substituted for CaCl_2_ in the recording solution for asynchronous EPSC (asEPSC) recordings. NASPM (100 μM; Tocris), the selective antagonist for GluA2 subunit lacking AMPA receptors (Ca-permeable AMPA receptors), was added to the recording solution in a subset of recordings to verify Ca-permeable AMPAR insertion. Recording electrodes (thin-wall glass, 1.5 OD/1.12 ID; WPI Instruments) were made using a Brown-Flaming model P-97 electrode puller (Sutter Instruments, San Rafael, CA) to yield resistances between 3-6 MΩ. Electrodes were filled with either a potassium based solution (in mM): 135 KMeSO_4_, 12 NaCl, 0.5 EGTA, 10 HEPES, 2 Mg-ATP, 0.3 Tris-GTP (for sEPSC and some LTD recordings), or a cesium based solution (in mM): 120 CsMeSO_4_, 15 CsCl, 8 NaCl, 10 HEPES, 0.2 EGTA, 10 TEA-Cl, 4 Mg-ATP, 0.3 Na-GTP, 0.1 spermine, and 5 QX-314-Cl (for, AMPA/NMDA ratio, asEPSC, RI, NASPM, and some LTD recordings). Chemicals were obtained from Sigma-Aldrich or Fisher Scientific unless otherwise noted.

### Data Acquisition and Analysis

Whole-cell voltage-clamp recordings were conducted on shNAc D1-MSNs surrounded by ChR2-expressing terminals identified by epifluorescent illumination of tdTomato and eYFP. Excitatory postsynaptic currents (EPSCs) were acquired using an amplifier (PC-One; Dagan) filtered at 1 kHz and digitized at 10-20 kHz with a Digidata 1440B interface board using pClamp 10.6 (Axon Instruments). A blue light LED (465 nm; Plexon) was coupled to the objective via a fiber coupler (IS-OGP/OLY, Siskiyou Corporation). Following the acquisition of the whole-cell configuration D1-MSNs were held at −80 mV and oEPSCs were evoked with 4 ms blue light pulses. Light intensity was adjusted for each recording to elicit oEPSCs in a range from 150-250 pA. In experiments for long term plasticity, oEPSCs were evoked every 15 seconds for at least 10 minutes. To induce LTD, a conditioning stimulus of 500 light pulses (4 ms) at 1 Hz was paired with continuous postsynaptic depolarization from −80 mV to −50 mV. oEPSCs were then monitored for 30 minutes after pairing. The magnitude of LTD was calculated by averaging normalized (to baseline) oEPSC values from 20-30 minutes post pairing and comparing that to the average normalized oEPSCs during the 10-minute baseline. Our criterion for the expression of plasticity required that the difference between averaged oEPSCs from the baseline and post pairing be greater than 2 standard deviations. Correlations between the amount of ethanol consumed (g/kg) in the last drinking session and the magnitude of LTD (% baseline) were tested using the non-parametic Spearman’s rank order test. Spontaneous EPSCs (sEPSCs) were recorded for 3 minutes separated into 20 consecutive sweeps 10-min after achieving whole-cell configuration. Events greater than 5 pA were included for analysis using Clampfit v10.6. Asynchronous EPSCs (asEPSCs) were evoked by blue light stimulation in the presence of SrCl_2_. Analysis of asEPSCs began 30ms after stimulus offset up to 400ms. Frequency and average amplitude of sEPSCs and asEPSCs were determined (using Clampfit Template Search). Paired-pulse ratios were acquired by applying two stimuli of equal intensity, separated by an interstimulus interval (ISI) of 50 ms and calculated as a ratio of oEPSC 2/oEPSC 1. NMDA/AMPA ratios were calculated as the ratio of the oEPSC recorded at +40 mV, 50 ms after afferent stimulation (NMDA) and the peak oEPSC recorded at –80 mV (AMPA). NMDAR currents were recorded at +40 mV and were pharmacologically isolated using 50 μM picrotoxin, 10 μM DNQX, and 1 μM CGP 52432. For rectification index recordings a current-voltage plot was generated measuring AMPAR-mediated currents at the following holding potentials (in mV): −80, −40, 0, +20, +40. The RI was determined as the ratio of oEPSC amplitudes at +40 and −80 mV. For all recordings, membrane and access resistances were monitored throughout the experiments. Cells in which access resistance varied more than 35% were not included in analysis. The maximum access resistance allowed for recordings was 35 MΩ. Data was analyzed using a one-way ANOVA and Bonferroni post hoc test or Student’s t test in which statistical significance from baseline for within each group was defined as p < 0.05. A repeated measures four-way ANOVA with Greenhouse-Geisser corrections for violations of sphericity and Bonferroni post hoc test or Student’s t test was used to analyze drinking data.

## RESULTS

### Ventral hippocampal glutamatergic input to shNAc is capable of expressing NMDAR-LTD

To study excitatory inputs from vHipp to D1-MSNs in shNAc, we injected AAV2-CaMKIIa-hChR2(H134R)-eYFP into the vHipp of male *Drd1a*-tdTomato mice. After 13 weeks, we detected ChR2-eYFP expression in vHipp neurons and in terminals present in the shNAc (Figure 1 A-B). Prior studies indicated that photo-activation of ChR2 in glutamatergic terminals is sufficient for eliciting EPSCs in MSNs as well as eliciting plasticity (Britt et al., 2012; Pascoli et al., 2014). Consistent with this, blue light (465 nm, 4 ms pulses) reliably evoked EPSCs in D1-MSNs (oEPSC) (Figure 1B). Previously, electrical stimulation could induce NMDAR-dependent LTD in D1-MSNs of the shNAc (Jeanes et al., 2014; Renteria et al., 2018). To examine possible LTD of excitatory vHipp inputs, we combined light stimulation (4 ms pulses at 1 Hz for 500 sweeps) with postsynaptic membrane depolarization to −50 mV. Five hundred second paring was sufficient to produce LTD of oEPSC amplitudes (Figure 1 C-D). Extracellular perfusion of the NMDAR antagonist APV (50 μM) before and during the optical pairing protocol prevented the expression of LTD (Figure 1 C-D) confirming this form of optically-induced plasticity is dependent on the activation of NMDA receptors.

**Figure 1.**
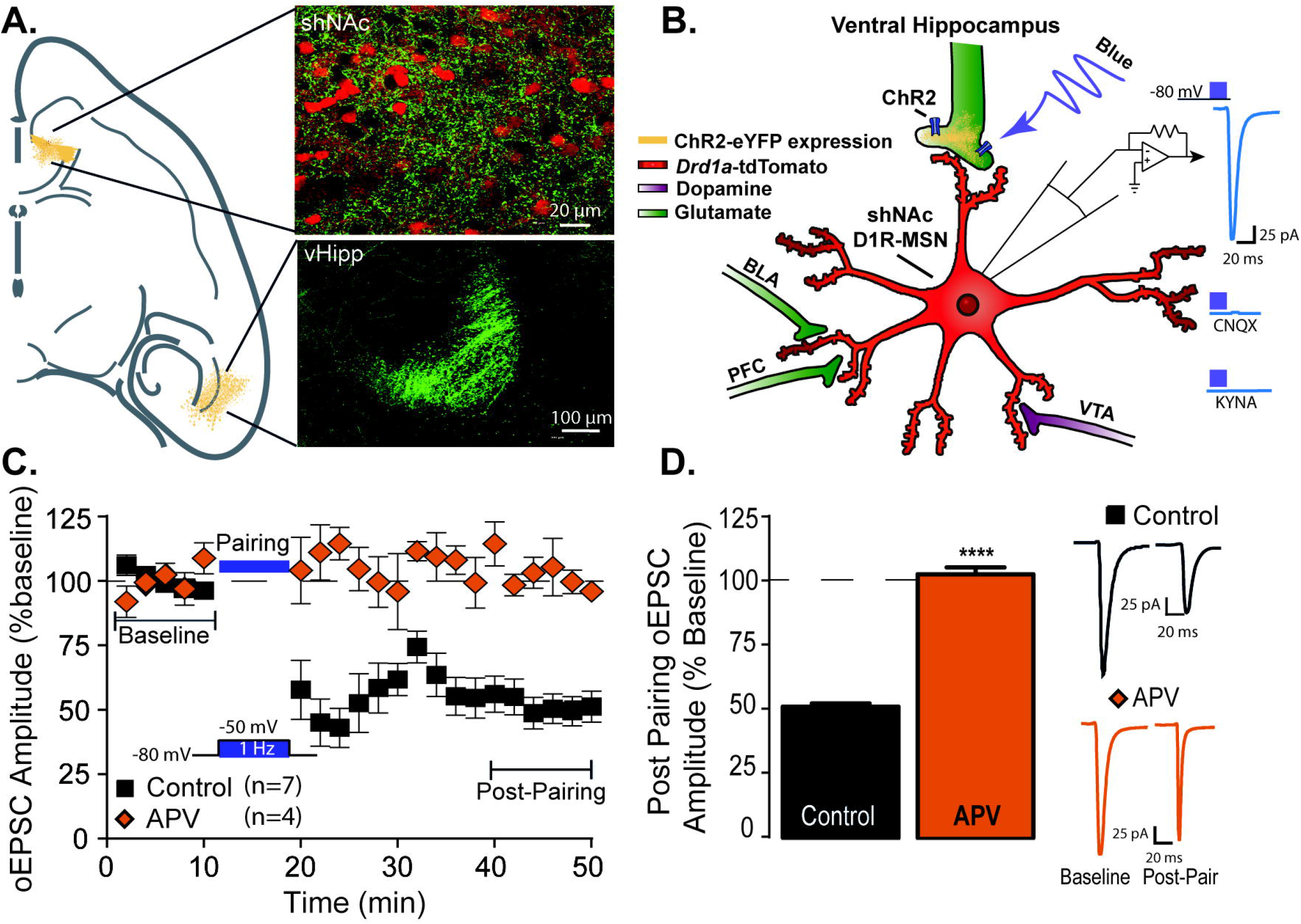
Optical low frequency stimulation paired with membrane depolarization of D1-MSNs induces NMDAR-LTD in vHipp-shNAc circuit. (A) Injection of AAV2-CaMKII-ChR2-eYFP into vHipp resulted in eYFP expression in vHipp glutamatergic neurons and in the terminals within the NAc shell. D1-MSNs were visualized by *tdTomato* fluorescence. (B) Whole cell patch clamp recordings were conducted in D1-MSNs and oEPSCs were elicited by blue light activation of ChR2 present in vHipp terminals within the NAc shell. oEPSCs are AMPA receptor dependent and were abolished in the presence of the AMPAR antagonist CNQX or non-selective antagonist KYNA. (C) A pairing protocol consisting of 1 Hz light stimulation (500 s) of vHipp terminals paired with postsynaptic membrane depolarization from −80 to −50 mV resulted in LTD of vHipp oEPSCs in D1-MSNs. The inclusion of the NMDAR antagonist D-APV in the bath prevented LTD expression. (D). Bar graphs representing average oEPSC amplitudes as a percent baseline ± S.E.M. during postpairing period (min 40-50). Student’s t-test: t(9)=6.3, ****P = 0.0001 APV vs control (Control: 51.19 ± 5.73 n=7 cells from 6 mice; APV: 102.6 ± 3.59, n= 4 cells from 4 mice).

### Acute *in vitro* ethanol exposure inhibits ventral hippocampal-accumbal glutamatergic plasticity

Acute ethanol can inhibit NMDA receptors (Martin et al., 1991; Thomas et al., 1998; Hendricson et al., 2004) as well as alter expression of LTD in the NAc (Jeanes et al., 2011, 2014, Renteria et al., 2017, 2018). We therefore examined the effect of acute ethanol on optically evoked LTD of the vHipp input to shNAc D1-MSNs. Bath application of a moderately to strongly intoxicating concentration of ethanol (40 mM), equivalent to ≈ 0.2 blood alcohol concentration (BAC), completely inhibited LTD expression (Figure 2A-B). Bath application of a low intoxicating concentration of ethanol (20 mM), equivalent to ≈ 0.1 BAC, also prevented induction of LTD (Figure 2A-B). We observed a significant difference in the average post-pairing oEPSC amplitude between both ethanol treatment groups and the ethanol naïve control group, with the ethanol naïve cells exhibiting robust reduction in oEPSC amplitude and little or no change in amplitude in the ethanol exposed cells (Figure 2D). A significant reduction in oEPSC amplitudes in the 20 mM exposed cells contrasts with our previous report using electrical stimulation, where we did not observe a disruption of LTD at this concentration of ethanol (Jeanes et al., 2011).

**Figure 2.**
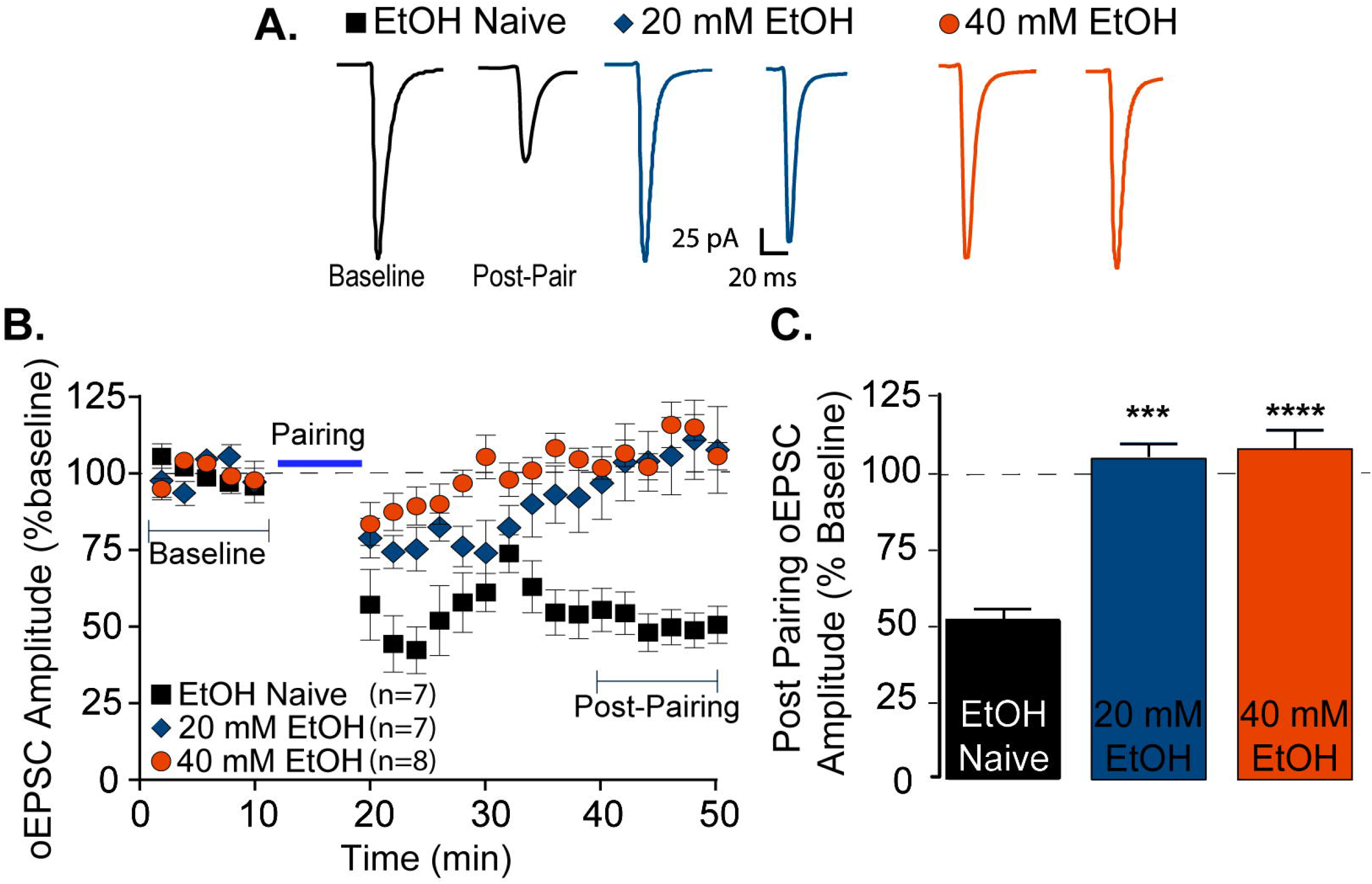
Acute *in vitro* ethanol exposure abolishes the expression of glutamatergic plasticity of vHipp-shNAc D1-MSNs. Escalating concentrations of ethanol applied *in vitro* in the recording bath disrupt the expression of NMDAR-LTD. (A) Sample traces of averaged baseline and post-pairing oEPSCs (40 sweeps, 10 min) of a single representative recording from each group. (B) A pairing protocol elicited LTD in ethanol naïve neurons. A total loss of LTD expression was observed during exposure to low (20 mM) and moderate (40 mM) concentrations of ethanol. (C) Bar graphs representing average oEPSC amplitudes as a percent baseline ± S.E.M. during post-pairing period (min 40-50). (EtOH Naïve: 51.19 ± 5.73, n=7 cells from 6 mice; 20 mM: 105 ± 11.47, n=7 cells from 4 mice; 40 mM: 108.1 ± 2.49, n=8 cells from 6 mice). ANOVA: F _(2, 19)_ = 19.25, P < 0.0001. Bonferroni *post hoc* tests: ***P < 0.001, 20 mM EtOH vs naïve; ****P < 0.0001, 40 mM EtOH vs naïve.

### CIE vapor exposure induces escalation of volitional ethanol intake in mice

Our *in vitro* findings suggest that the vHipp-shNAc circuit is sensitive to acute ethanol exposure. To examine the neural adaptations produced by *in vivo* ethanol exposure within this circuit, animals were given free access to consume ethanol and were then subjected to either ethanol vapor or air exposure. Two weeks after stereotaxic injection of the viral vector for ChR2-eYFP or control-eYFP, mice were subjected to daily, limited access, two bottle choice (2BC) drinking for 21 days prior to the first vapor (or air control) exposure (see Figure 3A for details). The last five days of drinking were measured as the baseline of consumption and the dose of ethanol consumed during the baseline was not different between the air and ethanol vapor exposed groups (Student’s t test: t(_6_) = 0.6835, P = 0.5198; Air: 1.51 ± 0.04, Vapor: 1.56 ± 0.07). CIE vapor exposure resulted in an escalation in ethanol consumption (g/kg) following each bout of chambering as compared to baseline intake. No escalation in ethanol consumption was observed in the air exposed mice. There was also no difference in intake between animals injected with ChR2-eYFP or eYFP (control) virus (Figure 3B). The CIE vapor induced increase in volitional ethanol consumption mirror those previously reported by our lab (Renteria et al., 2018).

**Figure 3.**
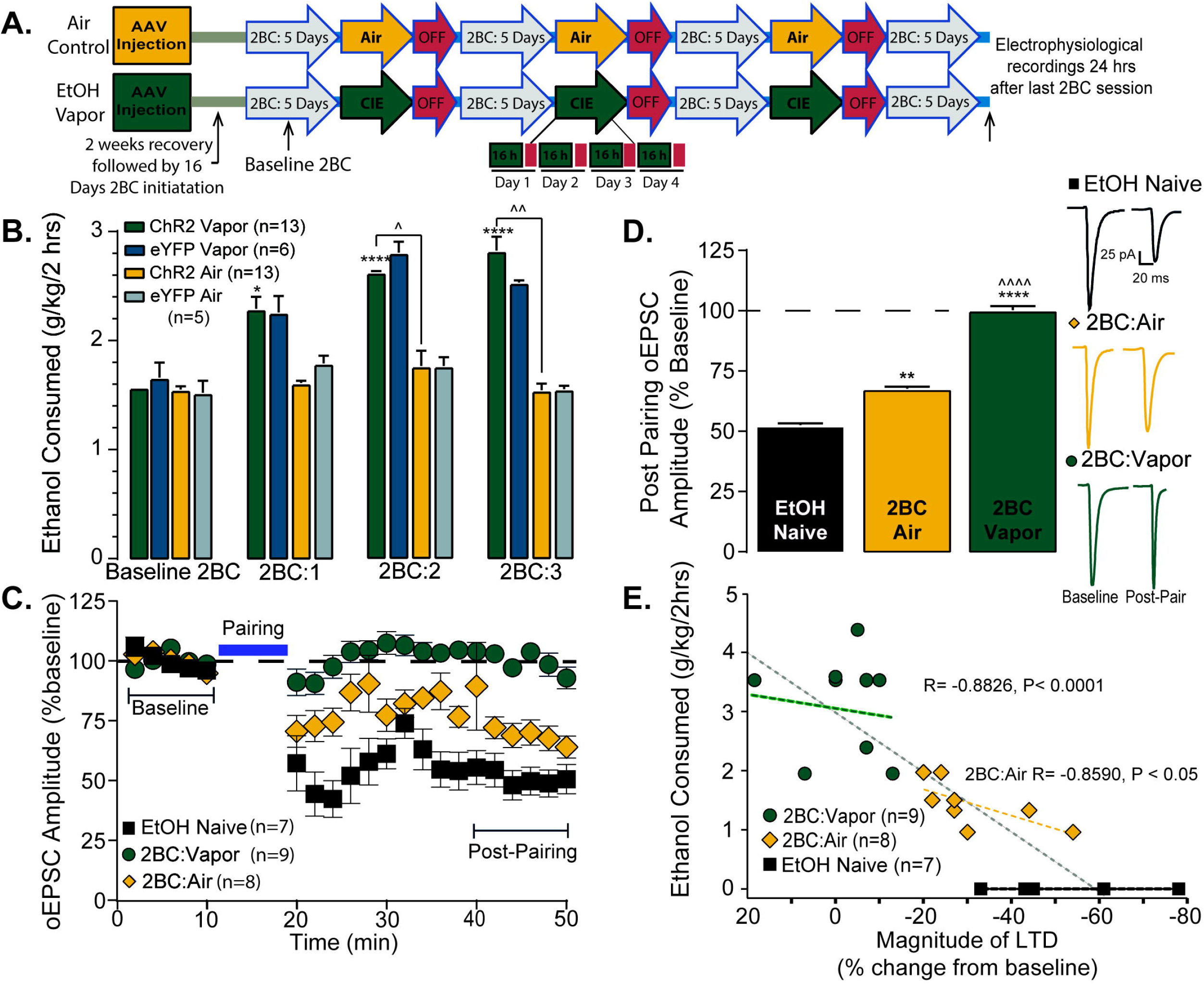
*In vivo* ethanol exposure disrupts glutamatergic plasticity of vHipp-shNAc D1-MSNs. (A) The CIE exposure consisted of 3 bouts of ethanol vapor (or air control) exposure separated by 3-day withdrawal and 5-day 2BC drinking (15% ethanol vs water) periods. Slices were prepared approximately 24 hours after the last ethanol drinking period. (B) Average consumption of ethanol (g/kg/2hrs) ± S.E.M. for each 2BC drinking period. Animals exposed to ethanol vapor exhibited an increase in 2BC drinking after each bout of vapor exposure. Four-way ANOVA (with Greenhouse-Geisser corrected degrees of freedom) indicated a significant main effect of drinking period (F_(2.389, 78.841)_ = 10.44 P <0.0001), with a significant treatment group by drinking session interaction (F_(2.389, 78.841)_ = 4.37, P = 0.011) and no virus group by drinking session interaction (F_(2389, 78.841)_ = 0.31, P = 0.772). *P < 0.05 vs baseline, ****P < 0.0001 vs baseline, ^P < 0.05 vs air, ^^P < 0.01 vs air. (C) The pairing protocol resulted in LTD in naïve animals, but 24 hours after the last drinking session failed to elicit LTD in the vapor exposed animals. Relative to ethanol naïve, air exposed animals exhibited a reduction in LTD magnitude 24 hours after the last drinking period. (D) Bar graphs representing average EPSC amplitudes as a percent baseline ± S.E.M. during post-pairing period (min 40-50) for each treatment group (EtOH Naïve: 51.19 ± 5.73, Air: 72.03 ± 2.54, Vapor: 99.86 ± 2.96. 1; ANOVA: F _(2 21)_ = 42.31, P < 0.0001. Bonferroni *post hoc* tests: **P < 0.01 vs naive, ****P < 0.0001 vs naive, ^^^^P < 0.0001 vs air; Naïve n=7 cells from 6 mice, Air n=8 cells from 4 mice, Vapor n=9 cells from 5 mice. (E) Inverse correlation between amount ethanol consumed during the last session and the magnitude of LTD observed in the air exposed animals (Spearman’s R= −0.8590, P < 0.05), and an overall negative correlation with LTD magnitude and ethanol exposure (Spearman’s R= −0.8826, P < 0.0001).

### Ethanol consumption and CIE induced escalation of intake disrupts ventral hippocampal-accumbal glutamatergic plasticity

Previous studies have shown that NMDAR-dependent plasticity is disrupted after *in vivo* drug or ethanol exposure (Brebner et al., 2005; Kasanetz et al., 2010; Pascoli et al., 2012; Abrahao et al., 2013). Here, we examined whether induction of LTD in D1-MSNs of the vHipp-shNAc circuit was affected by prior exposure to escalating ethanol intake.

In ethanol vapor treated mice, NMDAR-LTD in the vHipp-shNAc circuit did not develop with pairing of optical stimulation and post-synaptic depolarization (Figure 3C-D). In 2BC air-exposed mice, NMDAR-LTD was expressed but was reduced as compared to ethanol naïve control animals (Figure 3C-D). We also observed that the amount of ethanol consumed during the last 2-hour drinking session was correlated with the magnitude of LTD. Recordings from air exposed mice showed a significant inverse relationship between ethanol consumption and the magnitude of LTD. Interestingly, recordings from vapor exposed animals showed no correlation; however there was an overall correlation between the amount of ethanol consumed and the magnitude of LTD when Vapor, Air, and ethanol naïve mice were analyzed (Figure 3E). These findings suggest that the amount of ethanol consumed, in the absence of vapor exposure, opposes the expression of vHipp-shNAc NMDAR-LTD.

### Ethanol consumption alters ventral hippocampal-accumbal glutamatergic signaling

Previously, Renteria et al. (2018) showed ethanol consumption and CIE vapor exposure enhances spontaneous EPSC (sEPSC) frequency 24 hours after the last drinking session (Renteria et al., 2018). Here, we measured both the frequency and amplitude of sEPSCs. In ethanol vapor exposed mice, sEPSC frequency and amplitude were increased 24 hours after the last drinking session (Figure 4A-B).

**Figure 4.**
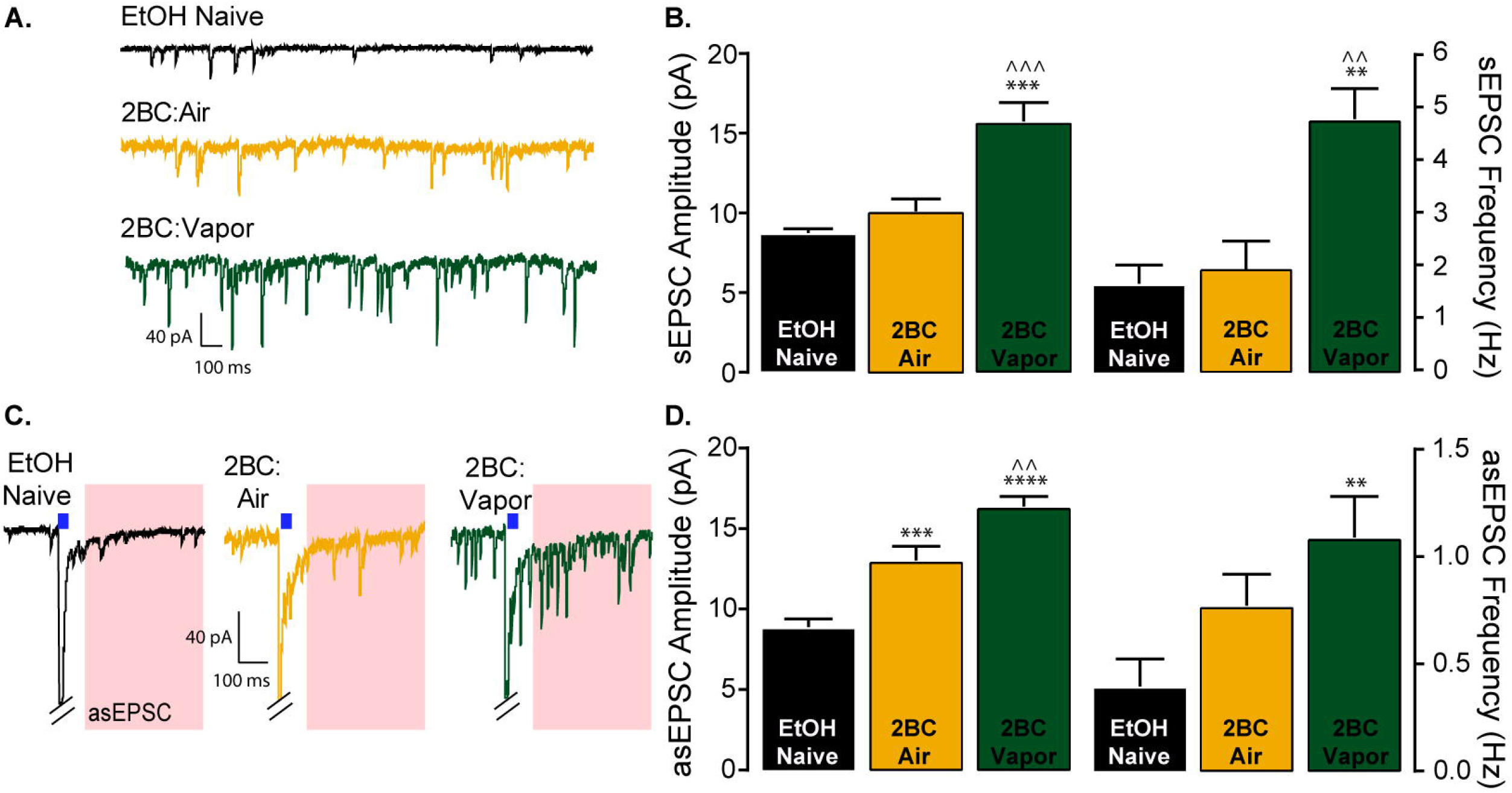
Ethanol experience enhances glutamatergic signaling in vHipp-shNAc circuit. (A) Representative traces of spontaneous EPSCs. (B) sEPSC amplitude (ANOVA: F _(2, 14)_ = 39.28, P < 0.0001, Bonferroni *post hoc* tests: ***P < 0.001 vs naïve, ^^^P < 0.001 vs air) and frequency (ANOVA: F _(2, 14)_ = 24.15, P < 0.0001, Bonferroni *post hoc* tests: **P < 0.01 vs naïve, ^^ P < 0.01 vs air) were elevated only in vapor exposed animals (Naïve n=5 cells from 5 mice; Air n=6 cells from 4 mice; Vapor n=6 cells from 4 mice). (C) Representative traces of asEPSCs; light stimulation in the presence of strontium elicit quantal like asynchronous events for a brief duration only from vHipp glutamatergic input. Red regions represent the time window where asEPSCs are observed (30-400 ms after light pulse). (D) Ethanol experience increased asEPSC amplitude (ANOVA: F _(2, 12)_ = 38.85, P < 0.0001, Bonferroni *post hoc* tests: ***P < 0.001 vs naïve, ****P < 0.0001 vs naïve, ^^P < 0.01 vs air). An increase in asEPSC frequency was only observed in vapor exposed animals (ANOVA: F _(2, 12)_ = 7.98, P < 0.01, Bonferroni *post hoc* test: **P < 0.01 vs naïve; n = 5 cells from 5 mice for all groups).

To examine circuit specific adaptations in glutamatergic signaling, we took advantage of ChR2 expression in vHipp terminals and used blue light stimulation to evoke quantal like events from vHipp terminals in the presence of strontium (asEPSC). Replacing calcium with strontium in the extracellular solution prolongs release events and results in quantal like release events occurring for a short duration following stimulation (analysis began 30ms after stimulus offset and ended at 400ms) (Gerdeman and Lovinger, 2001). Circuit specific analysis of glutamatergic signaling revealed a significant increase in asEPSC frequency in the vapor treated group compared to naïve animals (Figure 4C-D) suggesting that vapor exposure and an escalation in ethanol consumption leads to a presynaptic alteration in glutamatergic signaling. We noted a significant effect of ethanol consumption on asEPSC amplitude. Post hoc analysis indicates a significant increase in asEPSC amplitude in the air treated group as compared to naïve, as well as significant increase in asEPSC amplitude in the vapor exposed group as compared to naïve and air exposed animals (Figure 4C-D). These findings suggest that ethanol consumption is enhancing postsynaptic glutamatergic function at D1-MSNs synapses with the vHipp, with the largest enhancement in asEPSCs occurring following escalation in consumption produced by CIE vapor exposure.

Considering the observed alterations in asEPSC amplitude and frequency we attempted to recapitulate the presynaptic effect by measuring paired-pulse ratios (PPR) (2 pulses, 50ms interstimulus interval), which are inversely correlated with neurotransmitter release. We observed that ethanol consumption produced a decrease in the PPR (oEPSC 2 / oEPSC 1) (Figure 5A-B). Post hoc analysis indicated a significant reduction in PPR in both air and vapor treated groups, relative to ethanol naïve, suggesting that ethanol consumption leads to an increase in the probability of glutamate release from vHipp terminals onto D1-MSNs of the shNAc. Taken together, these findings indicate that ethanol consumption enhances glutamatergic signaling in the vHipp-shNAc circuit by reducing LTD and increasing pre-synaptic release.

**Figure 5.**
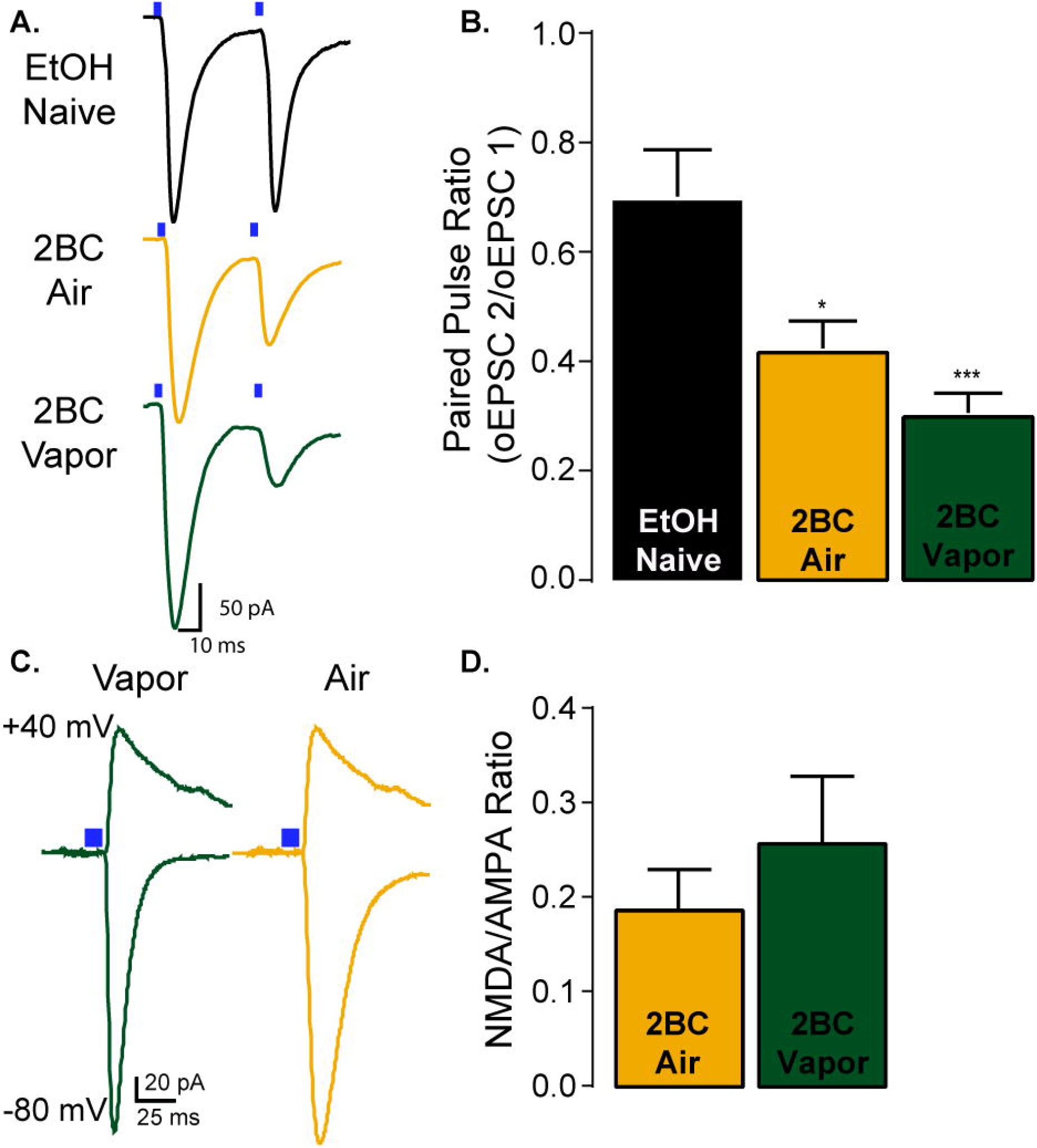
Ethanol experience enhances presynaptic glutamatergic signaling but not NMDAR function. (A) Representative traces of paired pulse (50 ms ISI) recordings. (B) Bar graphs representing paired pulse ratio (second oEPSC amplitude / first oEPSC amplitude). Ethanol experience reduced the paired pulse ratio with the maximum reduction in PPR observed in the vapor exposed group (ANOVA: F _(2, 21)_ = 13.67, ***P = 0.0002; Bonferroni *post hoc* tests: *P < 0.05 air vs naïve, ***P < 0.001 vapor vs naïve; Naïve n=5 cells from 3 mice; Air n=10 cells from 6 mice; Vapor n=9 cells from 5 mice). (C) Representative traces of NMDAR/AMPAR ratio recordings. AMPAR component was elicited at −80 mV holding potential. NMDAR component was elicited at +40 holding potential and measured at 50 ms after stimulus offset to allow for decay of any AMPAR mediated current. (D) Bar graphs representing the NMDAR/AMPAR ratio (NMDAR mediated EPSC amplitude / AMPAR mediated EPSC amplitude). No difference was observed in this ratio between the two groups; Student’s t-test: t(17)=0.7689 P = 0.45; Air 0.186 ± 0.04, Vapor 0.25 ± 0.07; Air n=8 cells from 6 mice; Vapor 11 cells from 6 mice).

### Ethanol consumption does not alter ventral hippocampal-accumbal NMDA receptor activity

Previous work demonstrated that NMDA receptor activity is enhanced twenty four hours after CIE vapor exposure as measured by electrically-evoked NMDA/AMPA ratios exhibiting enhanced NMDA amplitudes (Renteria et al., 2017). Here we hypothesized that NMDA receptor activity could be enhanced in the vHipp-shNAc circuit following the escalation of drinking produced by CIE vapor exposure. In the vapor or air treated mice, the amplitudes of NMDA/AMPA ratios in D1-MSNs were not significantly different (Figure 5 C-D). The absolute values of the NMDA/AMPA ratios for both groups were consistent with those previously shown in animals that did not undergo 2BC drinking and received air (control, naïve) exposure in a single 4-day bout of CIE treatment (Renteria et al., 2017).

### Ethanol consumption leads to the insertion of Ca-permeable AMPARs into ventral hippocampus-accumbal D1-MSN synapses

Insertion of Ca-permeable, rectifying, GluA2-lacking AMPARs occurs following exposure to and withdrawal from psychostimulants (Cull-Candy et al., 2006; Conrad et al., 2008), opiates (Hearing et al., 2016), and ethanol (38). Here we examined the potential change in rectification of vHipp-shNAc AMPARs following ethanol consumption. D1-MSNs prepared from mice 24 hours after 2BC drinking exhibited a rectification of AMPAR oEPSCs at positive holding potentials and an overall decrease in the rectification index (oEPSC amplitude at +40 mV/ oEPSC amplitude at −80 mV) (Figure 6A-B). To confirm that the alteration in rectification index in ethanol-experienced mice was due to the insertion of Ca-permeable AMPARs, we conducted light evoked current recordings and bath applied the GluA2 subunit-lacking AMPAR selective antagonist NASPM. We observed a significant reduction in oEPSC amplitudes in the presence of NASPM (Fig 6C-D) in both air and vapor exposed mice. Taken together these results indicate that the reduction in the rectification index observed after ethanol consumption is due to the insertion of Ca-permeable AMPARs.

**Figure 6.**
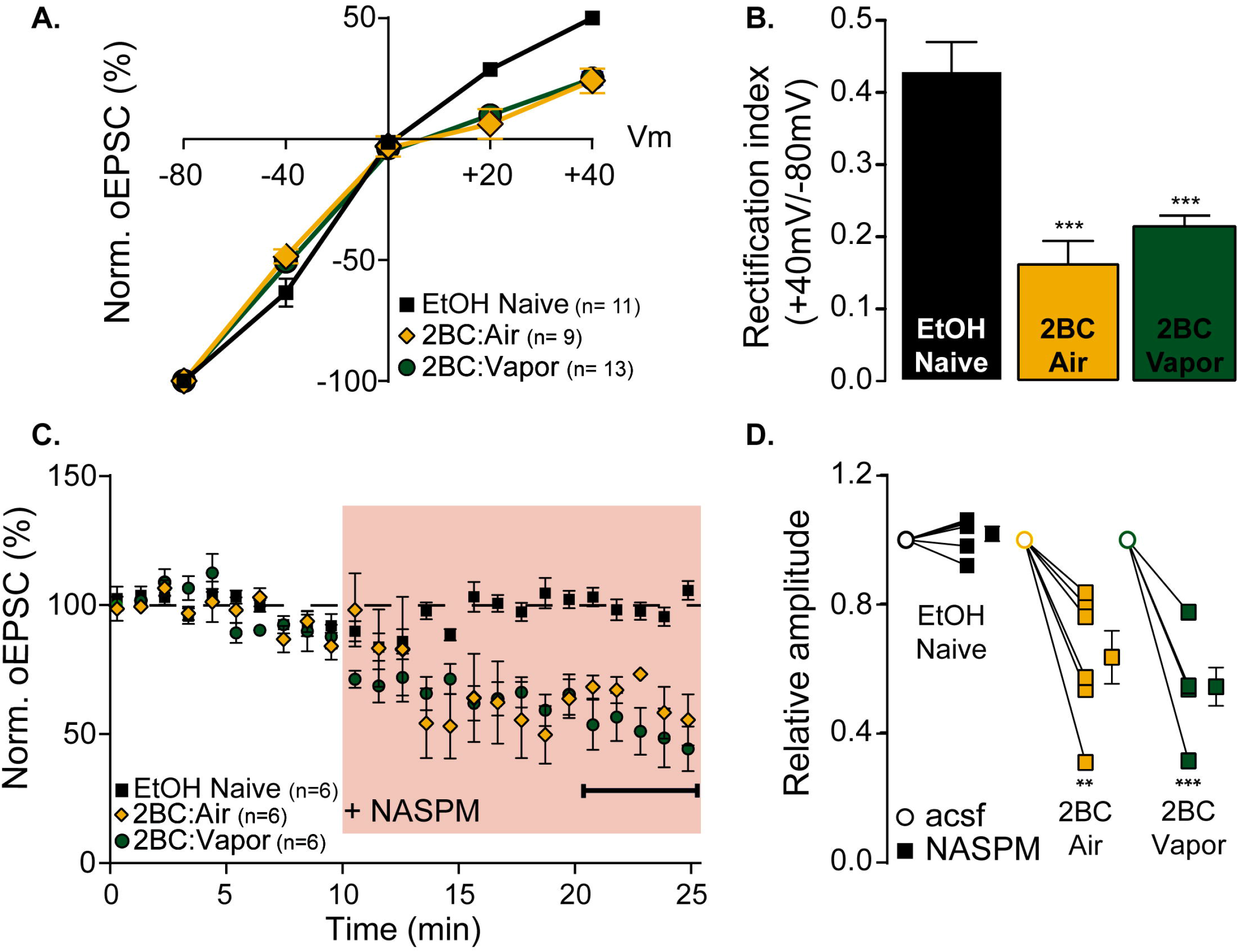
Ethanol experience results in the insertion of Ca-permeable AMPARs into vHipp shNAc synapses. (A) Current voltage (IV) relationship of AMPAR mediated EPSCs (in the presence of D-APV) in vHipp-shNAc D1-MSNs. Twenty-four hours after drinking a significant inward rectification is observed in ethanol experienced animals but not naïve animals (Naïve n= 11cells from 5 mice; Air n=9 cells from 6 mice; Vapor n=13 cells from 7 animals). (B) Rectification index (EPSC amplitude at +40 mV / EPSC amplitude at −80 mV) is decreased in both vapor and air exposed groups 24 hours after drinking (ANOVA: F _(2, 30)_ = 20.99, P < 0.0001; Bonferroni *post hoc* test ***P < 0.001 Air or Vapor vs Naïve). (C) Bath application of the Ca-permeable AMPAR selective antagonist NASPM produced a reduction in EPSC amplitudes in both air and vapor exposed animals (D) Averaged relative amplitudes of EPSCs before and after washing on NASPM. (ANOVA: F _(2, 15)_ = 17.51, P < 0.0001; Bonferroni *post hoc* tests: **P < 0.01 vs naïve, ***P < 0.001 vs naïve; mean relative amplitude ± S.E.M following NASPM (average of final 5 min): EtOH Naïve: 1.02 ± 0.02, Air: 0.64 ± 0.08, Vapor: 0.54 ± 0.06; n=6 cells from 6 animals for all groups).

## DISCUSSION

Here, we used pathway-specific optogenetics to focus exclusively on plasticity in the ventral hippocampal-accumbal circuitry in response to low and high alcohol drinking. We discovered that alcohol exposure enhanced excitatory synaptic transmission by reducing LTD, enhancing presynaptic release activity, and upregulating post-synaptic Ca-permeable AMPARs, with many of these enhancements present in both the low alcohol drinking (2BC: Air) and the high alcohol drinking (2BC: CIE) group. We discuss these results in the context of previous findings and the possible impact on the reward pathway.

In the present study, we focused on D1-MSNs in the shNAc to determine input specific neural adaptations produced by *in vitro* and *in vivo* ethanol exposure. Because vHipp input to the shNAc is the most robust glutamatergic input to this region (Britt et al., 2012), we examined this vHipp to shNAc circuit. We first verified that there is NMDAR-dependent LTD of optically evoked EPSCs, and then determined if plasticity of this isolated input onto D1-MSNs of the shNAc is altered by ethanol exposure. The present results confirm and extend prior work in several ways. A previous study, in which local electrical stimulation was used to evoke excitatory responses onto D1-MSNs in the shNAc, demonstrated that escalated drinking following CIE is accompanied by a total loss of LTD (Renteria et al., 2018). Similarly, we found that LTD was absent in vHipp-shNAc D1-MSN glutamatergic synapses following CIE. Furthermore, by including an ethanol naïve group, our experiments extend prior findings and indicate that 2BC drinking, which did not escalate in the Air group, alone can be sufficient to disrupt LTD – at least in the vHipp-shNAc pathway. In addition to vHipp-shNAc metaplasticity following ethanol intake, pre- and post-synaptic adaptations in glutamate signaling from the vHipp were observed. Increases in asEPSC frequency and reductions in the PPR suggest enhancements in release from the vHipp following drinking which are further enhanced by vapor exposure. Postsynaptic signaling alterations, identified by increases in asEPSC amplitude and putative changes in AMPAR subunit expression, also were present in the ethanol-experienced groups. However, no change in NMDA/AMPA ratio was observed between Air or Vapor exposed animals. Prior work suggests that vapor exposure enhances NMDAR activity in D1-MSNs (Renteria et al., 2017) but these recordings did not differentiate between presynaptic inputs. The lack of a change in the ratio could indicate that there is no change in NMDAR activity within the selected synaptic population (vHipp-shNAc). This result seems unlikely since enhancements in AMPAR activity were observed in asynchronous recordings. Its possible that an enhancement in NMDAR activity is masked by an accompanying enhancement in AMAPR activity resulting in no change to the ratio. Another interpretation would be that there is an enhancement in NMDAR activity in both the Air and Vapor groups and a comparison with control would be needed to confirm. We observed evidence of the insertion of GluA2-lacking, Ca-permeable, AMPARs into vHipp-shNAc synapses following ethanol consumption, indicated by a decrease in RI and a reduction in oEPSC amplitude following application of the selective antagonist NASPM. Taken together, these findings indicate that chronic ethanol experience promotes vHipp glutamatergic transmission to D1-MSNs in the shNAc.

It should be noted that the ethanol naïve mice that were used for comparison with the *in vivo* ethanol exposed animals were not handling controls; these mice did not receive injections, nor did they undergo chambering or tail blood collection. Thus, we cannot rule out the possibility that the differences between the ethanol naïve and the ethanol-experienced groups could be related to other aspects of the experimental handling. Nonetheless, we argue that the data are consistent with ethanol exposure being the relevant factor underlying the observed enhancements in glutamatergic transmission. For example, if experimental handling – which was essentially identical between the 2BC Air and 2BC Vapor groups – was the primary cause of glutamatergic adaptations, rather than ethanol exposure, then we would expect to observe no differences between these two groups. Yet, for nearly every adaptation, the Air group mean was an intermediate value between the ethanol naïve and Vapor group means – a pattern that is consistent with group differences being related to the amount of ethanol exposure. Only the measures indicative of increases in Ca-permeable AMPARs did not exhibit this pattern (both the Air and the Vapor group showed marked rectification at +40 mV and sensitivity to NASPM). However, even a single exposure to ethanol can be sufficient to promote an increase in GluA2-lacking AMPARs at glutamatergic synapses on shNAc D1-MSNs (Beckley et al., 2016). Thus, we propose that the lack of an intermediate effect on AMPAR composition in the Air group could be attributable to the high sensitivity of this phenomenon to ethanol.

Moreover, it is of particular interest that, in the Air group, ethanol consumption showed a strong negative correlation with the observed magnitude of LTD, suggesting that in higher drinking animals there is a corresponding adaptation in plasticity within D1-MSNs of the vHipp-shNAc circuit that shifts these neurons away from the ability to express synaptic depression. This observation is consistent with a prior report of a similar relationship between ethanol consumption and LTD magnitude in shNAc D1-MSNs after operant self-administration experience (Mangieri et al., 2017). We did not find a correlation between ethanol consumption and the magnitude of LTD in the vapor-treated group. Indeed, CIE vapor exposure appeared to abolish the ability to elicit LTD in vHipp-shNAc synapses on D1-MSNs, suggesting a shift in the threshold for the expression of LTP versus LTD (Jeanes et al., 2014). Prior work has shown that a metaplastic shift, from synaptic depression to synaptic potentiation, is present in glutamatergic shNAc D1-MSNs 24 hours after mice are passively exposed to high concentrations of ethanol using CIE vapor exposure (Jeanes et al., 2014; Renteria et al., 2017). Thus, the additional experience of ethanol vapor exposure likely accounts for the lack of a strong relationship between volitional consumption and plasticity in the Vapor group. Although there was not a significant correlation between plasticity and consumption within the Vapor group, plotting the extent of LTD (or LTP) for all three groups (Naïve, Air, and Vapor) as a function of ethanol consumption revealed a strong, linear relationship between ethanol exposure and plasticity of vHipp-shNAc glutamatergic synapses. In sum, these findings show that chronic exposure to increasing quantities of ethanol suppresses synaptic attenuation and enhances vHipp excitatory synaptic input to shNAc D1-MSNs in a manner dependent on the level of exposure.

In both 2BC:Air and 2BC:Vapor mice, we demonstrate an increase Ca-permeable AMPARs relative to ethanol naïve mice, which is likely produced by ethanol consumption. In many brain regions AMPAR activity is largely or solely mediated through GluA2 subunit-containing AMPARs, with some cells having basal Ca-permeable AMPAR activity (Conrad et al., 2008; Reimers et al., 2011; McGee et al., 2015). The subunit composition of Ca-permeable AMPARs confers faster onset and offset kinetics as well as substantially increasing the single-channel conductance. The calcium-permeability of these channels is also implicated to have critical functional consequences on the synapse following their expression (Liu and Cull-Candy, 2000; Henley and Wilkinson, 2013; Wu et al., 2017). Numerous types of drug exposure have been found to increase Ca-permeable AMPARs within the NAc, and predominantly within D1-MSNs. Ca-permeable AMPAR insertion in D1-MSNs has been reported following exposure to cocaine (Terrier et al., 2016) or opiates (Hearing et al., 2016; Russell et al., 2016), and following alcohol drinking (Beckley et al., 2016) or vapor exposure (Renteria et al., 2018). Enhancements in Ca-permeable AMPARs with the shNAc following alcohol (Beckley et al., 2016) and cocaine (James et al., 2014) exposure appear to be dependent on mTORC1 signaling. Furthermore, disrupting mTORC1 signaling in the NAc suppresses ethanol intake (Neasta et al., 2010) as well as cocaine seeking (James et al., 2014). Together, these findings suggest that insertion of Ca-permeable AMPARs in D1-MSNs of the shNAc is critical for drug related behaviors. Thus, drug-induced expression of Ca-permeable AMPARs may represent a common mechanism by which repeated drug exposure and withdrawal can regulate neuronal excitability and subsequent behavior.

Our findings of enhanced glutamatergic activity in ventral hippocampal to D1-MSN synapses in the low alcohol-consuming group provides a putative target for intervention that is present early in the process of the formation of dependence. Manipulations of neural transmission at this circuit during low alcohol exposure may abolish the increase in ethanol consumption following vapor exposure, thereby preventing escalation to a pathological state. This provides a potentially critical window for early intervention. If the adaptations in accumbal glutamatergic signaling are indeed critical for the expression of enhanced ethanol intake, pharmacological or physiological manipulations during the withdrawal periods following vapor exposure should prevent the increase in ethanol intake or even reduce the amount of ethanol consumed. For example, blocking Ca-permeable AMPAR activity in the NAc is one possible pharmacological approach to reducing ethanol consumption, which could be tested by infusing the Ca-permeable AMPAR antagonist NASPM directly into the shNAc. This hypothesis is supported by the finding that cocaine seeking is significantly reduced after infusing NASPM into the NAc (Schmidt et al., 2015). Another potential intervention during this window is activation of mGluR1 metabotropic glutamate receptors. Activation of mGluRs has been shown to reduce cue-induced responding for cocaine (McCutcheon et al., 2011b, 2011a), and these receptors are involved in ethanol drinking behaviors (Cozzoli et al., 2012, 2014). mGluR1 receptors can be engaged by stimulating glutamatergic terminals in the NAc at 13 Hz, eliciting mGluR1-dependent LTD resulting in the removal of Ca-permeable AMPARs. mGluR1-LTD has been observed to reverse cocaine-induced expression of Ca-permeable AMPARs and abolish cue-induced cocaine seeking (Pascoli et al., 2014). If this approach were to successfully reverse the expression of Ca-permeable AMPARs and abolish cue-induced ethanol seeking, it would further indicate mGluR1 as a promising candidate for pharmacological intervention for alcohol dependence and alcohol use disorders.

Here we have employed optogenetic tools to demonstrate alterations in glutamatergic signaling from the ventral hippocampus to the nucleus accumbens shell following ethanol consumption and the escalation in ethanol intake following CIE vapor exposure. Our findings provide further evidence that the insertion of Ca-permeable AMPA receptors following repeated drug exposure – documented by many other groups – also occurs following repeated alcohol consumption and highlight this phenomenon as a target for intervention.

## ACKNOWLEDGMENTS

Thank you to Daniela Carrizales for her assistance with animal behavior. Thank you to Dr. Paul Slesinger for his critical review of the manuscript. Funding from the National Institute on Alcohol Abuse and Alcoholism (AA015167 and AA016651).

